# Comparison of methods that use whole genome data to estimate the heritability and genetic architecture of complex traits

**DOI:** 10.1101/115527

**Authors:** Luke M. Evans, Rasool Tahmasbi, Scott I. Vrieze, Gonçalo R. Abecasis, Sayantan Das, Doug W. Bjelland, Teresa R. deCandia, Haplotype Reference Consortium, Michael E. Goddard, Benjamin M. Neale, Jian Yang, Peter M. Visscher, Matthew C. Keller

## Abstract

Heritability, *h^2^*, is a foundational concept in genetics, critical to understanding the genetic basis of complex traits. Recently-developed methods that estimate heritability from genotyped SNPs, *h^2^ _SNP_*, explain substantially more genetic variance than genome-wide significant loci, but less than classical estimates from twins and families. However, *h^2^_SNP_* estimates have yet to be comprehensively compared under a range of genetic architectures, making it difficult to draw conclusions from sometimes conflicting published estimates. Here, we used thousands of real whole genome sequences to simulate realistic phenotypes under a variety of genetic architectures, including those from very rare causal variants. We compared the performance of ten methods across different types of genotypic data (commercial SNP array positions, whole genome sequence variants, and imputed variants) and under differing causal variant frequencies, levels of stratification, and relatedness thresholds. These results provide guidance in interpreting past results and choosing optimal approaches for future studies. We then chose two methods (GREML-MS and GREML-LDMS) that best estimated overall *h^2^_SNP_* and the causal variant frequency spectra to six phenotypes in the UK Biobank using imputed genome-wide variants. Our results suggest that as imputation reference panels become larger and more diverse, estimates of the frequency distribution of causal variants will become increasingly unbiased and the vast majority of trait narrow-sense heritability will be accounted for.

## INTRODUCTION

Narrow-sense heritability, *h*^2^, the proportion of the total phenotypic variance due to additive genetic variation, is a fundamental concept of medical and quantitative genetics. In addition to providing an understanding of the genetic basis of traits, *h*^2^ determines the response to selection, the potential utility of individual genetic risk and trait prediction, and how much of the phenotypic variability could theoretically be accounted for in genome-wide association studies (GWAS)^1,2^. Importantly, while GWAS have now identified thousands of variants associated with complex traits^3–5^, the loci identified by these studies have typically explained only a small fraction of traits’ total heritability, with the remaining genetic variance termed “missing heritability.” This remaining unaccounted for genetic variance may be attributable to a variety of causes, including the role of (typically rare) variants poorly tagged by arrays, small effect common variants that do not reach genome-wide significance due to insufficient sample sizes, or inflated family-based *h*^2^ estimates^1, 6–8^.

While traditional family-based estimates of heritability, *h^2^_FAM_*, have provided valuable insights^9^, the use of close relatives means that estimates of additive genetic variance can be biased by factors shared by close relatives—for example, the joint action of non-additive genetic and common environmental effects can inflate estimates of additive genetic variation^10,11^. Recently-developed approaches that utilize unrelated individuals to estimate the variance explained by all genotyped single nucleotide polymorphisms (SNPs), denoted as *h^2^_SNP_*, have the advantage of being unaffected by these sources of bias, and for many traits have found that a large proportion of the heritability is captured by common variants^6,12,13^. For certain complex traits, such as height, little unexplained additive genetic variance remains, as *h^2^_SNP_* approaches *h*^2^_*FAM*_^7,12^. Despite this, *h*^2^_SNP_ estimates for most traits are still below *h^2^_FAM_*, with BMI a typical example where *h^2^_SNP_* ~0.27 while *h^2^_FAM_* ~0.4-0.6 (ref. ^12^). Thus, for many complex traits, including disease traits, much of the heritability remains unaccounted for.

A second application of these approaches is to better understand the genetic architecture of complex traits. Genetic architecture refers to the number, frequencies, effect sizes, and locations of causal variants (CVs) underlying trait variation. Methods for estimating heritability from SNPs have found that estimated genetic variance is proportional to chromosome length for numerous complex traits, including height, BMI, schizophrenia, depression, and metabolic traits, consistent with the hypothesis that these traits are influenced by hundreds to thousands of variants with small effects spread throughout the genome^5,6,8,12-16^. More recently, these methods have allowed insight into the frequency distribution and functional annotation of causal variants by partitioning SNPs into MAF bins and annotation categories^17,18^. Such methods have allowed insight into gene networks involved in complex traits^19^, and helped determine optimal strategies for large-scale genotyping, such as whether genotyped SNPs on commercial arrays with subsequent imputation can capture the genetic variation from all frequency classes of causal variants or if whole genome sequences instead are needed^12^.

A variety of methods to estimate *h*^2^_SNP_ and partition the genetic variance among sets of markers have been developed for these purposes. Many of these methods use one or more genetic relatedness matrices (GRMs) to estimate variances using restricted maximum likelihood (GREML)^6,12,17,20^. Manipulations of the GRM via treelet covariance smoothing^21^ or weighting by linkage disequilibrium (LD) tagging of SNPs^13^ have also been proposed. A much different approach, LD-score regression, estimates *h*^2^_*SNP*_ from GWAS summary statistics^22^. The performance of these methods has typically been evaluated via simulation by assuming that causal variants have the same properties, on average, as common SNPs found on commercial genotyping arrays. However, such an approach is problematic because SNPs are specifically selected because they are common, have unusually high LD with untyped SNPs, or have been implicated in disease (e.g., the Affymetrix Axiom chip used in the UK Biobank^23^). SNPs on arrays are therefore probably not reflective of typical CVs across the genome, and thus the ability of these methods to estimate *h*^2^_SNP_ or determine the genetic architecture of complex traits has not yet been properly assessed, nor have these methods been directly compared across conditions, such as levels of stratification or environmental confounding, that can cause biases. In particular, how the various methods perform with traits derived from very rare CVs may be quite different than how they perform on traits derived from common, welktagged CVs, such as those used on SNP arrays.

Here, we utilize thousands of recently-sequenced whole genomes to simulate complex phenotypes to test the performance of the most widely used SNP heritability estimation methods. We examine each method’s ability to estimate *h*^2^_*SNP*_ while varying the amount of population stratification, the frequency distributions of causal variants, and the type of whole-genome data analyzed (SNP array, imputed, and sequence). By using real sequence data to simulate phenotypes, the genotypic data we use are highly realistic with respect to LD, allele frequency distributions (with minor allele frequencies down to 3×10^−4^), variant density, and other genomic properties found in real data. Finally, we use the best-performing methods to estimate *h*^2^_SNP_ and examine genetic architecture for six complex traits using the UK Biobank. While *h*^2^_*SNP*_ estimation following imputation can account for the majority of the heritability, larger sample sizes and reference panels, or novel methods, will be needed to fully account for all the additive genetic variance in complex traits involving very rare causal variants.

## MATERIALS AND METHODS

### Samples and Population Structure

We simulated continuous phenotypes derived from whole genome sequence (WGS) data in the Haplotype Reference Consortium (HRC) dataset. Full details of the HRC can be found in McCarthy et al.^24^. Briefly, this resource comprises roughly 32,500 individual whole genome sequences from multiple whole-genome sequencing studies, with phased genotype calls available at all sites with a minor allele count of at least 5. The HRC contains world-wide populations, but the majority are of European (EUR) origin. This large collection allowed us to simulate phenotypes with differing genomic architectures under realistic patters of LD structure, stratification, and relatedness with the whole genomes. We obtained permission to access the following HRC cohorts (recruitment region & sample size): AMD (Europe & worldwidee 3,189), BIPOLAR (European ancestrye 2,487), GECCO (European ancestrye 1,112), GOT2D (Europe, 2,709), HUNT (Norwaye 1,023), SARDINIA (Sardiniae 3,445), TWINS (Minnesotae 1,325), 1000 Genomes (worldwidee 2,495), UK10K (UKe 3,715) (see web resources for HRC information including specific cohorts). The subset of the HRC data we accessed totaled 21,500 whole genome sequences comprising 38,913,048 biallelic SNPs.

Our goal was to assess the bias and precision of various *h^2^_SNP_* estimation methods using data similar to that typically used in GWAS and *h*^2^_*SNP*_ analyses. In order to mimic this kind of data, we first extracted variant positions corresponding to a widely-used commercially available genotyping array, the UKBiobank Affymetrix Axiom array. We performed principal components analysis using flashpca^25^ on 133,603 SNPs after LD and MAF pruning (plink2^26^ commands-maf 0.05—indep-pairwise 1000 400 0.2), extracting the first ten PCs, and performing K-means clustering in R^27^. We used the 1000 Genomes individuals in the HRC as anchor points for ancestry and identified 19,478 individuals of European descent, including individuals of Finnish and Sardinian ancestry (Figure S1).

To identify subsets of these 19,478 individuals spanning different levels of genetic heterogeneity, we reran PCA with only these individuals, then proceeded to identify four increasingly homogenous subgroups within them using K-means clustering (Fig. 1). The most stratified group contained all EUR samples (N=19,478). The somewhat stratified group excluded Sardinian and Finnish samples (N=14,424). The low stratification group contained only northern/western European samples (N=11,243), and the least stratified (homogeneous) group was a subset of British ancestry samples (N=8,506). We used GCTA^20^ to estimate relatedness and remove samples so that the maximum relatedness was 0.1 within each of the four samples. In the most homogeneous (smallest) sample, this left 8,201 individuals. To avoid confounding sample size with degree of stratification, we randomly chose 8,201 of the unrelated individuals from within each of the other three more stratified subsamples. Our purpose in identifying these groups was to vary the amount of genetic heterogeneity within a sample, similar to what might be found across a range of different GWAS samples, rather than formal population assignment or classification of individuals. We also identified individuals with relatedness less than 0.05 within each group, and used both subsets to examine how a 0.1 or 0.05 relatedness cutoff influences *h^2^_SNP_* estimates. Sample sizes when using the 0.05 relatedness cutoff were 7792, 8115, 8129, and 8186 for the four genetic structure subsamples.

**Figure 1.**
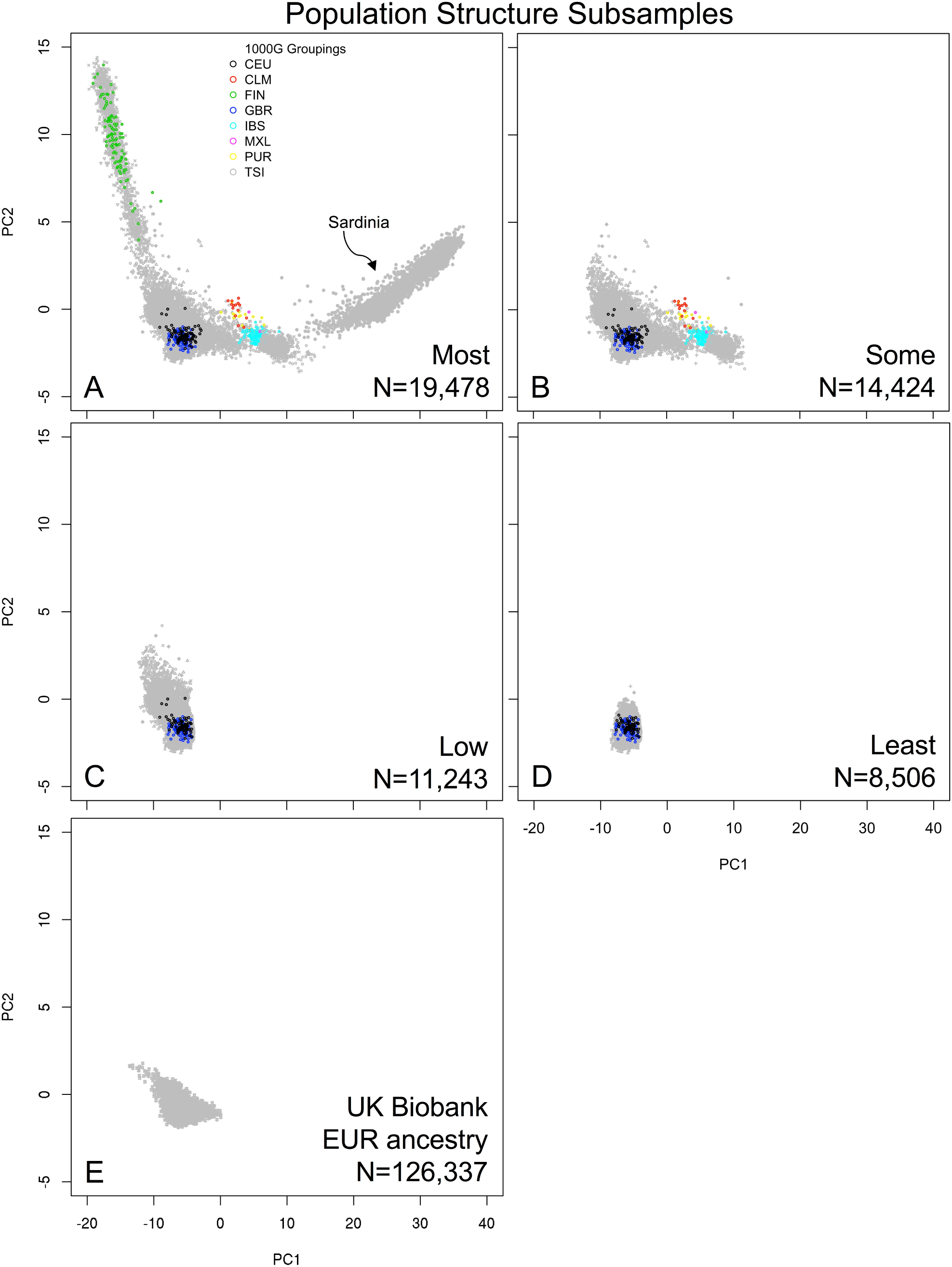
Population structure subsamples of European ancestry individuals in the HRC (A-D). and UK Biobank individuals projected onto these axes (E). Total sample sizes are shown in each panel. To keep sample size constant across stratification level, we randomly sampled 8,201 individuals with relatedness < 0.1 (the number of unrelated individuals in the most homogeneous and smallest set in panel D) from each subsample to create the subsamples used in the simulations.

### Simulated Phenotypes Using Whole Genome Sequencing Data

To assess how methods performed on a range of genetic architectures, we simulated phenotypes from CVs drawn randomly from five MAF ranges from the whole genome sequence data: common (MAF≥0.05), uncommon (0.01≤MAF<0.05), rare (0.0025≤MAF<0.01), very rare (0.0003≤MAF<0.0025), and all variants that had a minor allele count (MAC) of at least 5 (MAF≥0.0003) (Fig. S2). Phenotypes were generated from 1,000 CVs from the model *y_i_* = *g_i_ + e_i_*, where *gi* = Σ*w_ik_β_k_w_ik_, w_ik_* is the genotype (coded as 0, 1, or 2) of individual *i* at the *k^th^* CV, and *β_k_* is the *k^th^* allelic effect size, drawn from ~N(0,1/[2*p_k_*(1-*p_k_*)]), where *p_k_* is the MAF of allele *k* within a population subset. This model therefore assumes larger average additive effect sizes for rarer variants. The *g_i_*’s were standardized and added to residual error drawn from ~N(0, (1-*h^2^*)/*h^2^*) for a *h^2^* of 0.5 for simulated phenotypes. A total of 100 repetitions were simulated for phenotypes derived from each CV MAF range and for each of the four population stratification subsets. It is important to note that we did not simulate any phenotypic effects as a function of ancestry within any of the subsamples, and thus biases related to stratification in our results were due to the genotypic (e.g., long-range LD), not phenotypic, effects of stratification.

### SNPs, WGS, and Imputed Variants

Most marker heritability studies utilize commonly available commercial arrays, and estimates of *h^2^_SNP_* reflect how well SNPs on these arrays tag CVs. In particular, CVs with low MAF or that exist in regions of low LD are typically tagged poorly by SNP arrays^6,13^ and *h^2^_SNP_* < *h^2^* in these situations. Alternatively, as large WGS reference panels (e.g., 1KG, UK10K, HRC) become increasingly available, imputing genome-wide variants based on SNP arrays is an attractive option for capturing more and rarer genetic variants than possible on arrays, although imputation accuracy declines with MAF^12^. Finally, using WGS data to estimate GRMs should reflect relatedness at all CVs, including those that are rare or in low LD with other SNPs. Although WGS data in phenotyped samples is not yet widely available at the sample sizes required for precise estimation of *h^2^_SNP_*, we include it as a benchmark for results based on array and imputed data and because large WGS samples are likely to become increasingly available in the future. We therefore tested each of these data types (array, imputed, and WGS variants) using each of the methods described below to determine how much of the heritability can be captured from each data type, and how closely results from imputed data mimic those from WGS data.

From the HRC sequence data (the WGS dataset), we extracted positions corresponding to the Axiom array as noted above (the array SNP dataset) with MAF>0.01. To impute, we used the 8,201 unrelated individuals in each population stratification set and added their close relatives (relatedness > 0.1) back into the sample as described below in the GREML-SC method description. We added these close relatives back in to the target imputation set in order to a) remove close relatives from the reference panel which would artificially increase imputation accuracy, and b) because some of the methods described below require the use of closely related individuals. We phased these individuals using SHAPEIT2^28^, imputed using minimac3^29^, and retained variants with imputation *R*^2^≥0.3 (ref.^12^). We used the HRC sequence data as our imputation reference panel after removing all target (8201 unrelated + relatives) individuals, thereby assuring ~independence (no relatedness) between the target and reference panels.

Final reference panel sizes for the four structure subsamples were 11,584; 12,799; 12,785; and 12,994. Reducing the sample size of the reference panel likely resulted in poorer imputation than had we used the full HRC panel but was nevertheless substantially larger than reference panels used in most past imputation procedures (e.g., 1,000 Genomes). Moreover, because the target and reference samples were from the same populations and the same cohorts, the imputation quality is likely higher than most GWAS samples would obtain. However, given that the HRC has become a widely-used imputation reference panel, our imputation quality is probably roughly reflective of imputation quality using modern procedures.

The amount of tagging throughout the genome differs between the various commercial arrays^12^, and these differences may lead to differing *h^2^_SNP_* estimates. To assess this, for the GREML-SC and GREML-MS methods (see below) using array positions data, we compared results from the Axiom array to those from the Illumina Omni2.5 array. For reference, MAF distributions of the different data types for two of the structure subsamples are shown in Figure S2.

### Heritability Estimation Methods Tested

Numerous methods have recently been developed to estimate *h^2^_SNP_* and partition genetic variance using genomic data. Among these, we compared the most widely used, including the various single and multiple component GREML approaches implemented in the GCTA software^6,12,17^, approaches that specifically take into account how LD influences the tagging of nearby sites by SNPs, those that use related and unrelated samples to account for rare and common variant effects^8^, those that denoise the GRM using treelet covariance smoothing^21^, those that relate the effect sizes of SNPS from a GWAS to their degree of LD tagging ^19,22^, and computationally efficient mixed model approaches^18^. Here, we briefly describe our implementation of each of these methods; for additional information on the methods themselves, see the above references. For all methods except LD-Score Regression and BOLT-REML (described below), we generated GRMs following the procedures of each method, and estimated *h^2^_SNP_* using GCTA^20^. In all models, variance component estimates were unconstrained (e.g., by using the –reml-no-constrain option of GCTA), and included 20 PCs (10 from worldwide PCA and 10 from the specific subsample PCA) as continuous covariates and sequencing cohort as a categorical covariate.

### Single Component GREML (GREML-SC)

Yang et al.^6^ introduced the single component GRM approach using a mixed-effects model, with GRM entries:

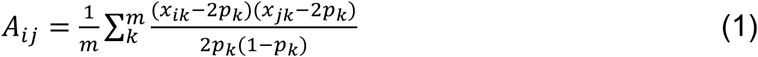

where *m* is the number of SNPs, *x_jk_* is the genotype (coded as 0, 1, or 2) of individual *j* at the *k*^th^ locus, and *p_k_* is the MAF of the *k*^th^ locus. The variance of the phenotypes is

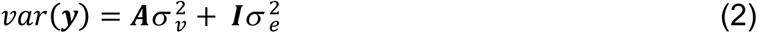

where the variance explained by the SNPs (*σ*^2^_*v*_) and error variance (*σ*^2^_*e*_) are estimated using restricted maximum likelihood (REML) implemented in the GCTA package^20^. The proportion of the total variance explained by all SNPs is then a measure of heritability (*h^2^_SNP_* = **σ*^2^_v_* / (**σ*^2^_v_ +*σ*^2^_e_*)). Typically, the set of *m* SNPs used to build the GRM is the set of SNPs with MAF≥0.01 (hereafter “common SNPs”) and unrelated individuals (relatedness ≤ 0.05). Because the Axiom array contains some rare markers, we compared this approach to one using all SNPs with MAC≥5 (hereafter “all SNPs”) in each particular stratification subsample, as well as to an approach using less stringent relatedness thresholds (relatedness < 0.10 and no relatedness threshold). For analyses that used no relatedness threshold, inclusion of close relatives increased our sample sizes to 9916, 8701, 8715, and 8506 for the samples with most, some, low, and least stratification, respectively (Fig. 1).

### MAF-Stratified GREML (GREML-MS)

Biased estimates of *h^2^_SNP_* are expected when using the GREML-SC method if the MAF distribution of the CVs does not match the MAF distribution of SNPs used to generate the GRM^17^. Stratifying variants into MAF classes and using a multiple GRM GREML approach can mitigate this bias and can also partition the genetic variance into that explained by different MAF categories of SNPs, lending insight into the genetic architecture of complex traits^12,30^. We applied this approach using 4 MAF categories, matching the CV MAF categories used for phenotype simulation.

### LD- and MAF-Stratified GREML (GREML-LDMS)

Extending the GREML-MS method to account for different levels of LD throughout the genome, Yang et al.^12^ introduced an LD score-stratified method to the GREML-MS approach. GREML-LDMS stratifies variants according to both MAF categories as well as an LD-score, defined as the sum of *r^2^* between the focal variant and all other variants in a window. We estimated LD scores using the default settings in GCTA (10Mb block size with a 5Mb overlap), and stratified variants into LD score quartiles. Combined with the four MAF categories above, we used 16 GRMs for this approach.

### Single Component and MAF-Stratified LD-Adjusted Kinships (LDAK-SC and LDAK-MS)

Speed et al.^13^ noted that because LD varies across the genome, CVs in regions of high LD are given disproportionate weight by eqn. (1) above. They proposed a method to weight SNPs according to local LD, which potentially corrects for the bias introduced when there is variation in how well CVs are tagged by SNPs. We used LDAK5^13^ to estimate these LD-weighted GRMs. This approach thins SNPs in very high LD first to reduce redundant tagging, then estimates SNP weights that are inversely proportional to their average LD with other SNPs. We also applied the MAF-stratified approach described above with the LDAK method (LDAK-MS). For the single component model (LDAK-SC), we used all SNPs (MAC≥5) as well as only common SNPs (MAF≥0.01) to build the GRM. For the MAF-stratified approach, following recommendations in the LDAK documentation, we estimated variant weights over the union of all variants (MAC≥5), then computed GRMs for each MAF class separately. We then applied the multiple GRM method with these LDAK-weighted GRMs to estimate *h^2^_SNP_* using GCTA.

### Extended Genealogy with Thresholded GRMs

Zaitlen et al^8^. introduced a method to simultaneously estimate the full narrow-sense heritability (incorporating the effects of poorly tagged SNPs) and *h^2^_SNP_* using two GRMs in a sample containing close relatives. The first GRM contains relatedness from SNPs for all individuals while relatedness estimates below a threshold, *t*, are set to 0 in the second GRM. The first GRM, therefore, contains information on allele sharing of (mostly common) variants in unrelated and related individuals and is used to estimate *h^2^_SNP_*, while the second only contains information from closely related individuals, presumably reflecting sharing of both common and rare CVs, and provides an estimate of what we call *h^2^_IBS>t_* (following Zaitlen et al.^8^). The sum of *h^2^_IBS>t_* and *h^2^_SNP_* should therefore provide an estimate of total *h^2^*, similar to *h^2^_FAM_.*, with all the same potential biases that exist in *h^2^_FAM_* estimates from designs that use close relatives. We tested two relatedness thresholds (*t* ≤ 0.05 and 0.1) for the second GRM. By necessity, all analyses using the relatedness thresholded GRM approach included close relatives.

### Treelet Covariance Smoothing (TCS)

Crossett et al.^21^ noted that the GRM estimates (particularly for unrelated individuals) are inherently noisy. They proposed a method to smooth the estimates using treelet covariance smoothing (TCS) to obtain more accurate estimates of relatedness. Their method takes advantage of the hierarchical nature of relatedness in samples to obtain better estimates of *A*_*ij*_ among unrelated individuals. We replicated their methods, using common SNPs (MAF≥0.01) and including related individuals, and implemented the TCS method in the *treelet* R package^31^. TCS requires identifying a smoothing parameter, *λ* (distinct from the genomic control inflation factor *λ*_GC_). Crossett et al.^21^ propose two methods to optimize *λ*, one based on minimizing the GREML likelihood and one based on minimizing a loss function (Η(*λ*)) at different levels of *λ* based on subsamples of the SNPs. With the large number of simulations across stratification subsamples and genetic architectures, minimizing the GREML likelihood for each simulated phenotype was not feasible. Minimizing H(*λ*) using the second approach requires estimating the GRM and applying the TCS method to over 50 subsets of data, also impractical computationally with over 8,000 individuals. We therefore used a modification of the 2^nd^ approach. We built GRMs from 2000 randomly chosen individuals from each stratification subsample and optimized *λ* for each subsample following the published methodology (Fig. S3), then applied the optimal *λ* to the full GRM of over 8,000 individuals.

### LD-Score Regression

LD-score regression uses a different approach to estimating *h^2^_SNP_*. Rather than estimating relatedness within a sample for use in mixed-model GREML analysis, LD-score regression regresses GWAS test statistics (*χ*^2^) on SNPs’ LD scores, which reflect the degree to which each SNP is correlated with surrounding SNPs ^19,22^. For a polygenic model, the expected GWAS test statistic of variant *j*, χ^2^_j_, is

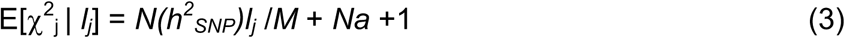

where *N* is the sample size, *M* is the number of SNPs, *l_j_* is the LD score (= Σ*_k_r^2^_jk_*) measuring the tagging of surrounding variants by SNP *j*, and *a* is a measure of confounding biases arising from stratification and cryptic relatedness. Thus, regressing GWAS test statistics on per-variant LD scores allows for both estimation of *h^2^_SNP_* and assessing the degree of confounding or polygenicity of a trait^22^. Bulik-Sullivan et al^22^ argue that LD-score regression provides unbiased estimates of *h^2^_SNP_* regardless of whether GWAS test statistics are estimated with or without controlling for ancestry or environmental covariates or relatedness. Here, we estimated GWAS test statistics using plink2 without controlling for ancestry covariates, controlling for ancestry covariates (20 PCs and sequencing cohort as above), and controlling for ancestry covariates as fixed effects in a mixed model that included a kinship matrix. For the latter, we applied the GCTA leave-one-chromosome-out (LOCO) approach^32^; because the GCTA-LOCO approach is computationally intensive, we ran only 20 repetitions of each phenotype rather than 100, and did so only for the array SNP dataset. We used the *Idsc* package with default parameters (see URLs) to perform LD score regression. We calculated LD scores for all variants using the whole genome sequence data, including common and rare variants. As recommended by Bulik-Sullivan et al., we used unrelated individuals (relatedness ≤ 0.05) and only common variants to perform the LD score regression itself, because the relationship between the GWAS &^2^ and LD-score is unclear for rare (MAF<.01) SNPs.

LD score regression can also be used to partition heritability among annotations^19^. We applied this approach using the four MAF categories described above. Because our MAF categories included very rare variants, for this MAF-stratified LD score regression, we used GWAS test statistics from all variants (MAF≥0.0003, using the--not-5-50 flag in the ldsc package) while controlling for covariates as above.

## BOLT-REML

Unlike other GREML approaches, BOLT-REML uses a Monte Carlo approximation of the gradient for the likelihood function to reduce computation time and memory requirements in variance component estimation^18^. When using whole genome sequence and imputed variant data with >14M variants (see below), time required by BOLT-REML, even when highly parallelized, was prohibitive for 100 repetitions of each combination of variables we tested, as it scales with MN^1,5^, where M is the number of markers and N is the number of samples (see Supplementary Table 1 of Loh et al.^18^ for computational performance). Note that GREML takes longer for a single sample due to the length of time to create the GRM; in our simulations with GCTA-style approaches, the GRM computation was done only once, and therefore was much faster when estimating heritability for many repetitions created from randomly-drawn CVs with a single GRM. We therefore only applied BOLT-REML to the array dataset. We applied the method with a single component using either all array positions or only common markers (MAF>0.01) as well as a MAF-stratified approach with the same four MAF partitions and same covariates described above.

### Confounding between relatedness and shared environments

Many of the methods we tested use unrelated individuals to avoid the assumption of no shared environmental effect among near relatives^6^. However, several, such as the extended genealogy with thresholding, require the use of near relatives^6^. This could lead to confounding between relatedness estimates and shared environmental effects within families or closely related individuals if shared environmental effects are not modeled^7,33^. Indeed, Zaitlen et al.^8^ argue that such shared environmental effects were the likely cause of higher *h^2^_FAM_* estimates among relatives who shared an environment through cohabitation (e.g., half-siblings) compared to equally related relatives that did not share a cohabitation environment (e.g., grand-parents and grand-children). We therefore assessed whether *h^2^_SNP_* and *h^2^_FAM_* estimates are biased for methods that use closely related individuals when extended shared environmental effects are present but unmodeled.

We first identified all groups of individuals connected by at least one pairwise relatedness value > 0.2 (“extended families”). Note that many of the pairwise relationships within these extended families were below 0.2. For example, spouses are typically unrelated but are nevertheless defined as being in the same family if their offspring are present, and cousins would be defined as being in the same family if their parents were present in the sample. We then simulated phenotypes with a shared extended family environmental effect that accounted for 10% of the variance (*c^2^*=0.1). Simulations were similar to those described above, with genotypic values exactly the same as above, but with shared effects for each family drawn from ~N(0, V_c_), where V_*c*_ = *c^2^*Vg/h^2^*, V_g_ is the variance of genetic values, and *c^2^* is the proportion of the phenotypic variance due to shared environments, and residual error added as ~N(0, (1-*h^2^-c*^2^)*V_g_/*h^2^*), for a simulated *h^2^*=0.5, *c^2^*=0.1, and *e^2^*=0.4. We applied GREML-SC, LD score regression, and extended genealogy with thresholded GRMs using common variants from array SNPs controlling for the same covariates as above and without modeling the shared environmental effect. This tested whether methods are robust to violations of the assumption of no shared environmental effects on the phenotype.

### Heritability of Complex Traits in the UK Biobank

We estimated *h^2^_SNP_* for six continuous phenotypes in the UK Biobank using the methods (GREML-MS and GREML-LDMS) that produced consistently unbiased estimates of *h^2^* and partitioned the genetic variance most accurately in the simulations above. The UK Biobank is a large, publicly available resource of ~500,000 UK adults, with deep phenotyping, family history, and genotype data^23^. The current release includes ~150,000 individuals, primarily of European ancestry, genotyped on the Affymetrix Axiom platform, phased using SHAPEIT2 and imputed to a combined 1000 Genomes and UK10K reference panel (N=6,285 individuals). The details of the official UK Biobank genotyping and imputation methods in the released data can be found at http://biobank.ctsu.ox.ac.uk/crystal/docs/genotyping_qc.pdf and http://biobank.ctsu.ox.ac.uk/crystal/docs/impute_ukb_v1.pdf (accessed 29 Feb. 2016). We excluded individuals with no genetic data and those whose self-reported and genetic sex conflicted (data fields f.31.0.0 and f.22001.0.0). Poor quality samples identified by the UK Biobank and Affymetrix were also removed (f.220010.0.0) as were UKBiLEVE poor-quality samples (f.22051.0.0), leaving a total of 151,661 individuals. To reduce population stratification, we included only individuals of European ancestry in our analyses. The UK Biobank identified self-reported “British” individuals as “Caucasian” based on grouping of individuals with CEU individuals in PCA (see UK Biobank documentation). To these individuals (f.22006.0.0), we added those who self-identified as “White,” “Irish,” or “Any other white background” whose PC scores on the first four axes (f.22009.0.1-4) were within the range of the UK Biobank-identified “Caucasian” individuals, resulting in 126,338 individuals. We projected the UK Biobank samples onto the HRC PCA axes using the loadings from the HRC EUR individuals, demonstrating that the UK Biobank individuals we used in the analyses below are similar to the least stratified or unstratified subsamples of the HRC we used (Fig. 1). To estimate the GRMs, we separately used directly genotyped Axiom array positions as well as imputed genome-wide variants with IMPUTE info score ≥0.3.

We estimated *h^2^_SNP_* for the following traits in the UK Biobank (field ID number): height (f.50.0.0), body mass index (BMI; f.21001.0.0), whole-body impedance (f.23127.0.0), trunk fat percentage (f.23127.0.0), fluid intelligence (f.20016.0.0), and neuroticism (f.20127.0.0). We normalized phenotypes and removed observations greater than 5 standard deviations away from the mean. We included sex (f.31.0.0), UK Biobank assessment centre (f.54.0.0), genotype measurement batch (f.22000.0.0), and educational attainment (“qualification”, f.6138.0.0) as categorical covariates, and the Townsend deprivation index (f.189.0.0), age at assessment (f.21003.0.0), age at assessment squared, and the 15 PC scores from the UK Biobank (f.22009.0.1-15) as quantitative covariates.

For GREML-MS, we binned variants into eight MAF-categories: MAC≥5 & MAF<0.0001, 0.0001-0.001, 0.001-0.01, 0.01-0.1, 0.1-0.2, 0.2-0.3, 0.3-0.4, & 0.4-0.5. For GREML-LMDS, we were limited in the number of predictor GRMs to use due to computational constraints (1Tb of RAM); we therefore, used 4 MAF bins (common: MAF>0.05, uncommon: 0.01<MAF<0.05, rare: 0.0001<MAF<0.01, and very rare: MAC>5 & MAF<0.0001) and 2 LD-score bins (above and below the median LD-score).

## RESULTS

### Simulation Results

We found clear differences across methods, degree of stratification, and data types (array SNP, WGS, or imputed variants) in their ability to estimate the simulated *h^2^* for different CV MAF architectures (Figs. 2–3 and S4-S6, Tables S1-S3). Below, we describe results for each method in detail. Please refer to Figures 2–4, Figures S4-S6, and Tables S1-S5 for estimates of heritability, and Figures S7-S9 for estimates of the heritability standard errors.

**Figure 2.**
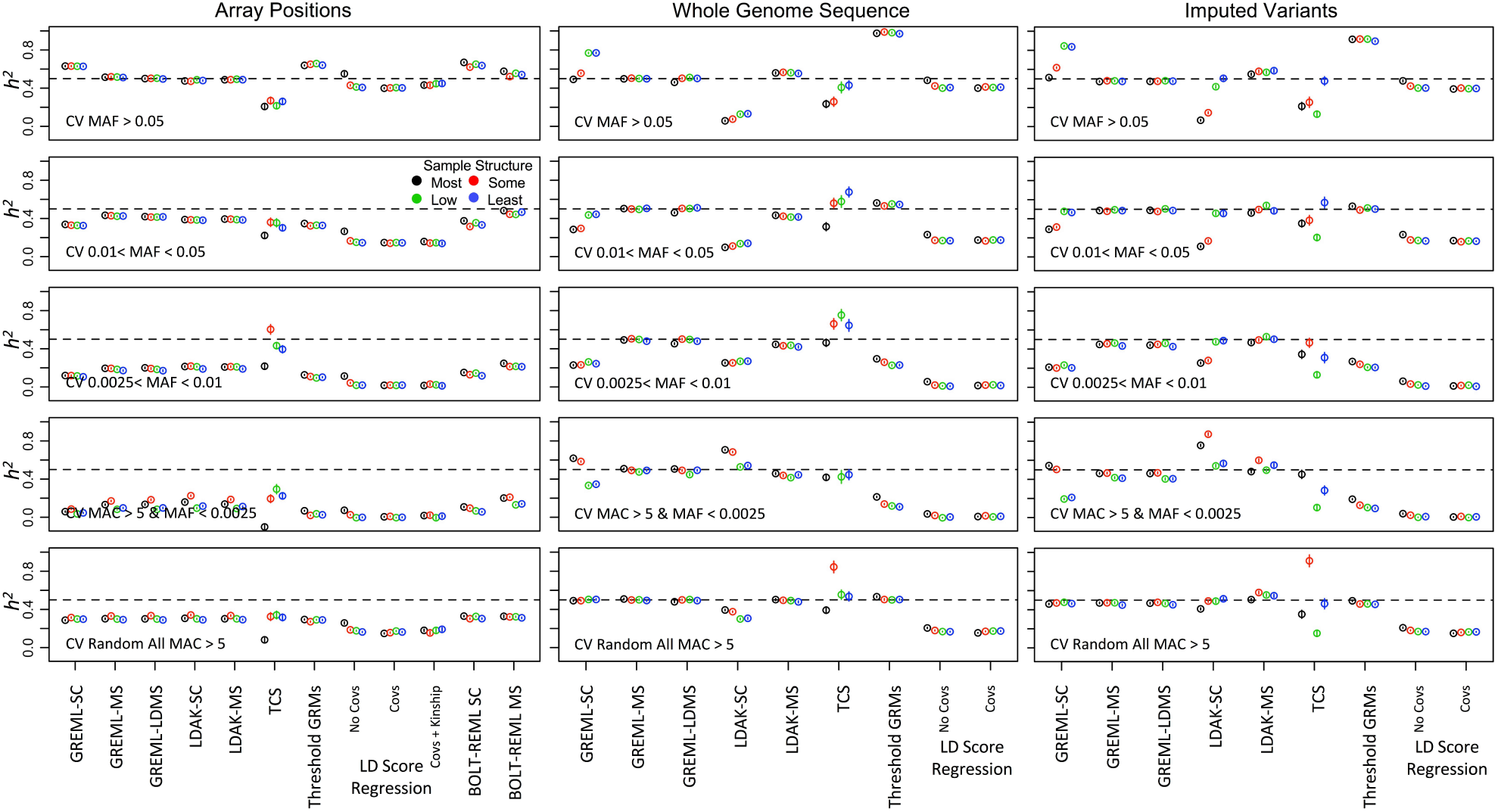
Average *h^2^_SNP_* estimates across 100 replicates (± SEM) from GRMs built from Axiom array positions (left), whole genome sequence data (center), or imputed genome-wide variants (right). Horizontal panels show MAF ranges (specified in insert) of 1,000 randomly chosen causal variants (CVs). Methods are listed on the X-axis as follows: Single component GREML (GREML-SC); MAF-stratified GREML (GREML-MS); LD-& MAF-stratified GREML (GREML-LDMS); Single-component Linkage Disequilibrium-Adjusted Kinships (LDAK-SC); MAF-stratified LDAK (LDAK-MS); Treelet Covariance Smoothing (TCS); Extended Genealogy with Thresholded GRMs; LD Score Regression using no PCs as covariates in GWAS, using PCs as covariates, or using both PCs and the kinship matrix; and Single Component and MAF-stratified BOLT-REML. Estimates are from samples of unrelated individuals (relatedness <0.05) except for samples used in the Threshold GRM method, which included all individuals. For the Threshold GRM method we plot *h^2^_SNP_* rather than total *h^2^* (*h^2^_SNP_* + *h^2^_IBS>t_*) from models where *t* = .05. Dotted line is the simulated (true) *h^2^* = 0.5. Colors represent the 4 subsamples varying in genetic structure. See Figs. S4-6 for estimates using different relatedness thresholds.

**Figure 3.**
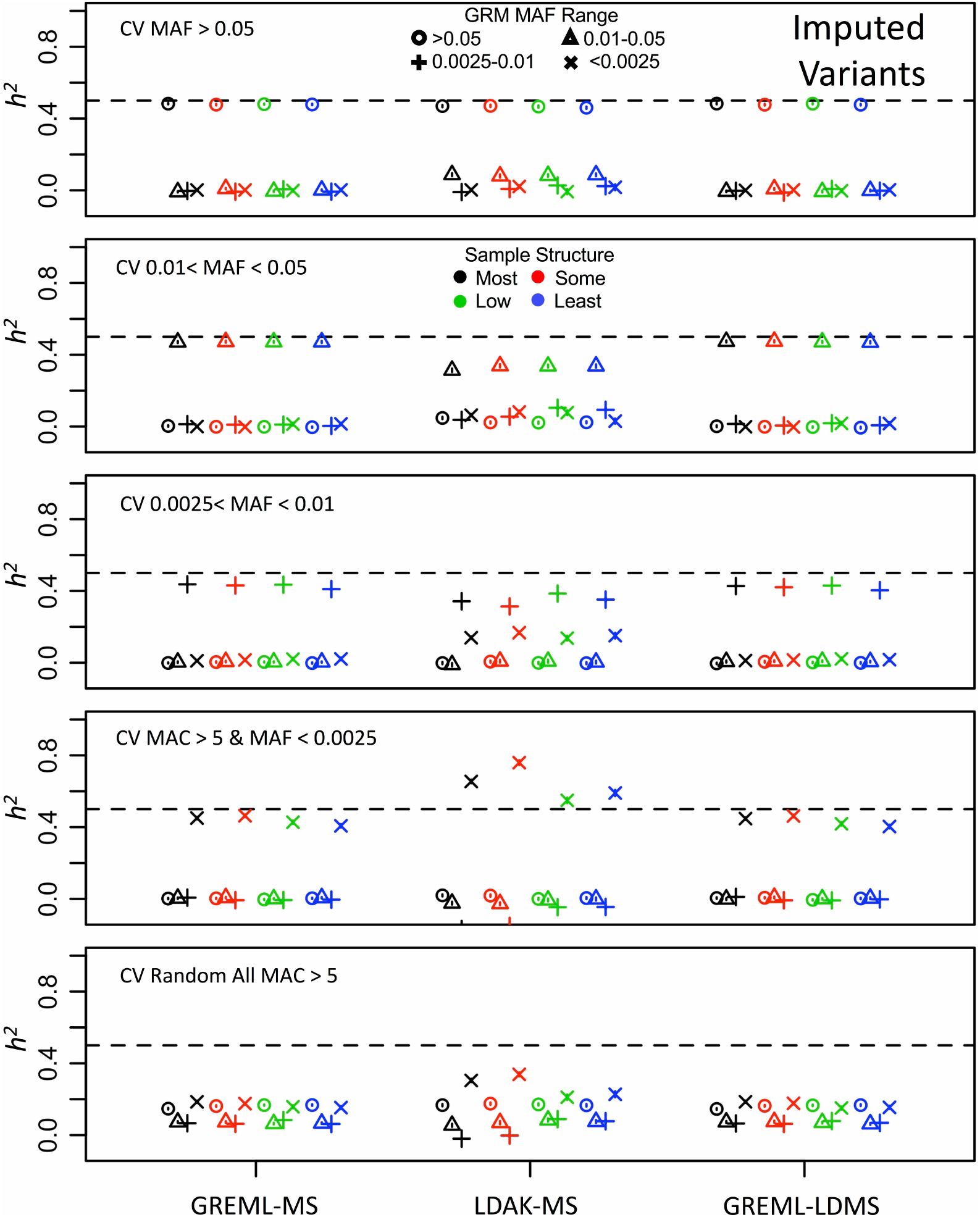
Average of 100 *h^2^_SNP_* estimates (± SEM) from GRMs constructed from imputed genome-wide variants of different MAF ranges (different symbols) in samples of unrelated (<0.05) individuals. Horizontal panels show MAF ranges (specified in insert) of 1,000 randomly chosen CVs and colors represent the 4 subsamples varying in genetic structure. GREML-MS & GREML-LDMS partition the phenotypic variance to the correct MAF-range GRM, while LDAK-MS often attributed genetic variance to incorrect GRMs.

**Figure 4.**
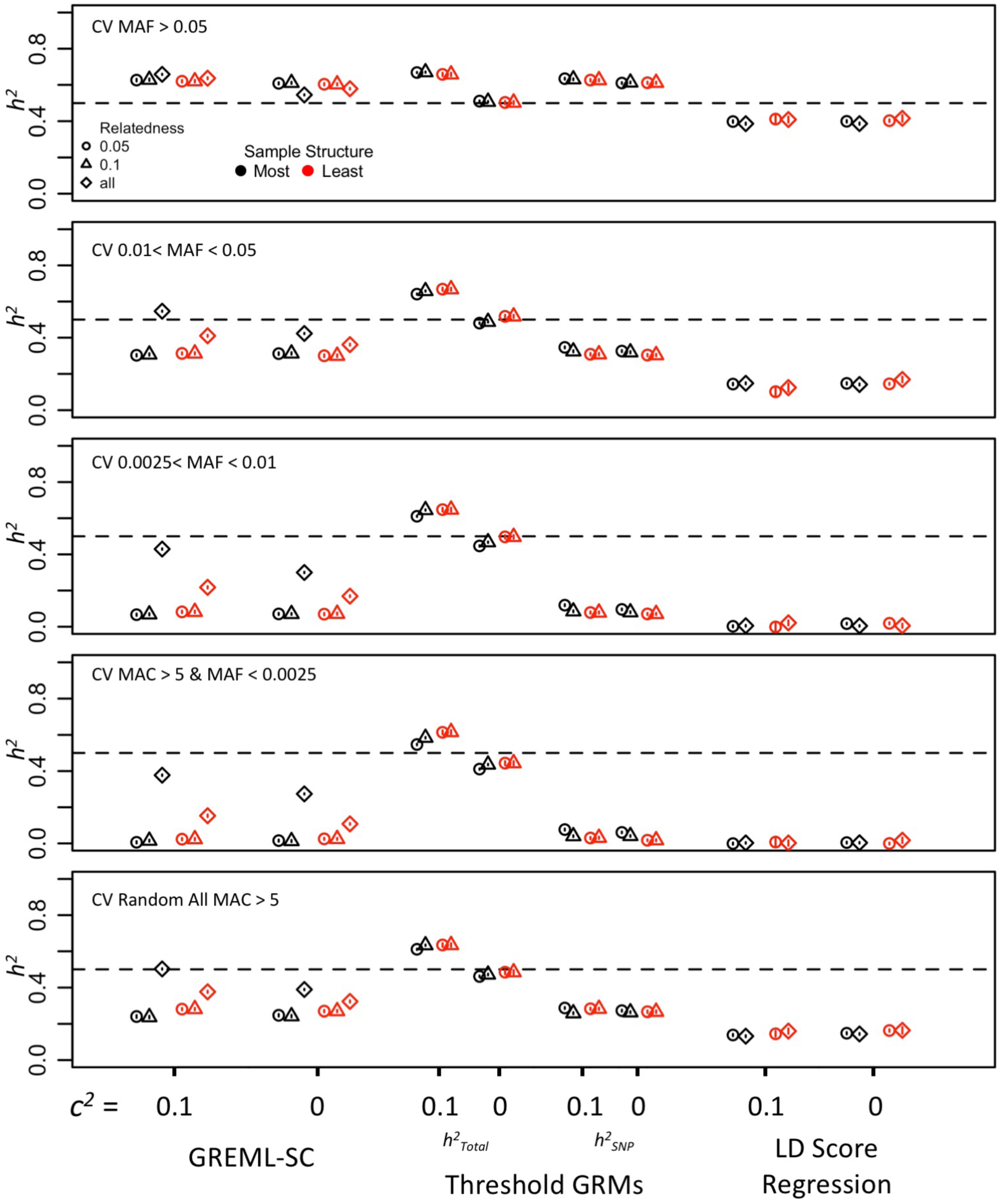
Mean heritability estimates (± SEM) from 100 replicates of phenotypes simulated with or without confounding shared environmental effects among families for three different methods (x axis) for different genetic architectures. GRMs were estimated using common (MAF>0.01) array SNP positions for the most structured and most homogeneous stratification subsamples only. Different symbols indicate the relatedness cutoffs used. For GREML-SC, we used three thresholds, including no relatedness cutoff (all individuals included). For LD Score Regression, we did not apply a 0.1 relatedness cutoff, as most studies will use a 0.05 or lower threshold for individuals included in GWAS. The threshold GRM approach requires all individuals, and the different symbols indicates the relatedness threshold (*t*) below which the thresholded GRM was set to 0. *h^2^_Total_* is the sum of both variance components, *h^2^_SNP_* is the variance component of the unthresholded GRM. Each horizontal panel indicates the minor allele frequency (MAF) range of the 1,000 randomly chosen causal variants (CV), with the range specified in the inset.

### Single Component GREML (GREML-SC)

Estimates of *h^2^_SNP_* using GREML-SC were highly sensitive to the CV allele frequencies, dataset type (SNPs, WGS, or imputed variants), level of stratification, and MAF cutoff for SNPs used to build the GRM. Using only Axiom array positions, *h^2^_SNP_* was overestimated by ~20% for common CV phenotypes, and progressively underestimated with rarer CVs, regardless of whether all or just common (MAF>0.01) SNPs were used to build the GRM. The underestimation of *h^2^_SNP_* when the GRM is built from SNPs that are more common on average than the CVs is well known^6^. It is due to a more general principle: when the average LD between CVs and the markers used to build the GRM is lower than the average LD among the markers themselves, *h^2^_SNP_* is underestimated^12^. Thus, *h^2^_SNP_* was underestimated for rare CVs because they tend to have lower LD with common markers than the common markers have with each other.

The overestimation of *h^2^_SNP_* for common CVs in our results is explained by the same principle-the average LD between CVs and markers is, in this case, higher than the average LD among markers used to build the GRM. This occurs for two reasons. First, the common CVs (MAF≥0.05) have higher MAF on average than the markers on the array (using either an MAF≥0.01 or MAC≥5 cutoff for the Axiom-based GRM computation). Second, markers on arrays are not chosen at random, but are typically chosen to minimally tag one another (to reduce redundancy) and to maximally tag variants not on the array, leading to lower average LD among markers than between markers and common variants not on the array (see also ref. ^13^). To understand if the overestimation of *h^2^_SNP_* for common CV phenotypes was unique to the Axiom array positions, we reran the analysis with SNPs on the Illumina Omni2.5 array and observed similar *h^2^_SNP_* inflations for common CVs on the Illumina array as well, although the impact of sample stratification appeared to more strongly influence the Illumina chip, perhaps due to the incorporation of a larger number of rare (MAF<0.01) variants on the Illumina array (Figs. S2 and S10).

Utilizing imputed or WGS data to build the GRMs resulted in complex patterns of *h^2^_SNP_* estimates depending on CV MAF class and stratification. Using a MAF>0.01 cutoff for imputed SNPs in building the GRM resulted in patterns similar to array-based estimates above (Fig. S5-S6), although the overestimates for common CVs were not as large, probably because the imputed markers used to build the GRM included all common SNPs rather than an overrepresentation of tag SNPs. On the other hand, when all imputed markers were used to build the GRM, GREML-SC estimates depended strongly on stratification level and the CV MAF. GREML-SC produced large overestimates for common CV phenotypes but underestimates for rarer CV phenotypes in unstratified samples (Fig. S6), as previously noted in Yang et al.^12^. The pattern was reversed for stratified samples: estimates of *h^2^_SNP_* were approximately unbiased for common CV phenotypes but underestimated for uncommon-to-rare CV phenotypes and overestimated for very rare CV phenotypes. Finally, when the frequency distribution of the CVs matched that of the WGS (e.g., randomly drawn from all WGS variants), the estimates were unbiased regardless of stratification when using WGS data to build the GRM (Fig. S5), but were slightly underestimated when using imputed data (Fig. S6), presumably due to imperfect imputation. The reason for this complex pattern of *h^2^_SNP_* estimates, where the effect of CV MAF depended on stratification, was likely due to changes in CV-marker and marker-marker LD as a function of stratification. The pattern of *h^2^_SNP_* estimates in unstratified samples is predictable based on the logic outlined above: when CVs are more common than the markers used to build the GRM, *h^2^_SNP_* is over-estimated, and vice-versa when CVs are less common than SNPs used to build the GRM. In highly stratified samples, however, very rare variants tend to be ancestry-specific and therefore weak proxies for variants elsewhere in the genome that predict ancestry (long-range LD). This makes the LD between very rare CVs and markers that predict ancestry elsewhere in the genome higher on average than the LD among the markers used to build the GRM, thereby inflating *h^2^_SNP_* estimates for very rare CV phenotypes in stratified samples.

These results underscore that *h^2^_SNP_* estimates from GREML-SC, the typical approach used, are sensitive to differences in average CV-marker LD vs. marker-marker LD (Fig. S11). This difference itself depends on complex interplays between the CV MAF distribution, the frequency distribution of markers used to build the GRM, and the level of stratification in the sample. Thus, *h^2^SNP* estimates using single-component GREML are highly context dependent, which may help explain the variation in estimates sometimes observed across studies for the same traits. Fortunately, stratifying SNPs based on MAF and LD, to which we turn next, largely ameliorates these issues.

### GREML using MAF-Stratified (GREML-MS) and LD- and MAF-Stratified (GREML-LDMS) GRMs

Genome partitioning using GREML-MS and GREML-LDMS produced *h^2^_SNP_* estimates that were substantially less biased and less sensitive to stratification than those from GREML-SC. GREML-MS *h^2^_SNP_* from array-based GRMs were underestimated for rarer CV phenotypes, as expected given the lack of LD between common array SNPs and rarer CVs, and were very slightly overestimated for common CV phenotypes, probably because of the LD properties of the SNPs chosen to be on the array (e.g., Illumina vs. Axiom positions, Fig. S10), as described in the previous section. GREML-MS using imputed variants slightly underestimated *h^2^_SNP_* for common to rare CV phenotypes. For very rare CV phenotypes, *h^2^_SNP_* was underestimated by ~18% in unstratified samples, likely due to poorer imputation quality for very rare SNPs, but underestimated by only ~7% in stratified samples. The higher estimates in stratified samples for very rare CVs is probably a lingering overestimation effect of long-range tagging of such variants in stratified samples. WGS-based estimates appeared unbiased for all combinations of CV MAF, relatedness, and stratification, with estimates all ~0.5.

Partitioning of the variance among the four MAF-stratified GRMs using GREML-MS allowed examination of the CV frequency distributions (Fig. S12, Table S4). GREML-MS estimated from GRMs built from array markers correctly apportioned the variation for common CV phenotypes, but as expected progressively underestimated *h^2^_SNP_* due to poor tagging of rare CVs with common SNPs (Fig. S12). Imputed variant GREML-MS provided more accurate estimates of the CV frequency distributions, but still underestimated the effects of rare and very rare CVs by as much as ~20% in unstratified samples (Fig. 3). Using WGS, the appropriate proportion of the variance explained by each MAF-stratified GRM in the model was recovered (Fig. S13, Table S4). Thus, the use of multiple GRMs based on MAF using imputed or WGS data produces generally accurate GREML estimates of both *h^2^sNP* and the CV frequency distribution, with only modest downward biases for very rare CVs when using imputed data.

The patterns of *h^2^_SNP_* estimates (Fig. 2–3, Figs. S12, S13) from GREML-LDMS were almost identical to those from GREML-MS, which might be expected because the CVs in our simulation were drawn at random within frequency bins and without regard to their LD. There were, however, two minor differences between the GREML-LDMS and GREML-MS results. First, for arrayabased GRMs (Fig. 2), estimates from GREML-LDMS for common CV phenotypes were unbiased, whereas those for GREML-MS were slightly overestimated. As noted above, array markers are more likely to tag common SNPs not on the array better than those on the array, leading to higher CV-marker than marker-marker LD and creating a slight upward bias. By binning by LD in addition to MAF, GREML-LDMS removes this source of bias, leading to unbiased *h^2^_SNP_* estimates for common CVs. Second, for unknown reasons, GREML-LDMS using whole genome sequence data gave slight (~3%) underestimates of *h^2^_SNP_* in highly stratified samples for rare to common CVs, but not for very rare CVs (Fig. 2 and S5). This effect was not apparent in imputed data, and may be simply sampling variance.

In summary, our findings suggest that using GREML-MS or GREML-LDMS on imputed data generally leads to accurate estimates of *h^2^_SNP_* and the CV allele frequency distributions, with only modest underestimation of variance due to rare and very rare CVs. Moreover, once large enough WGS datasets become available, the underestimation of rarer CVs should be largely ameliorated, although these methods can never estimate variance due to CVs that are so rare as to be unshared in a given sample.

### Single Component and MAF-Stratified LD-Adjusted Kinships (LDAK-SC and LDAK-MS)

Single component LD-adjusted estimates of the kinship matrix (LDAK-SC) downweights markers that better tag other SNPs, thereby correcting for the overestimation of *h^2^_SNP_* observed in GREML-SC for common CV phenotypes in array-based data due to redundant tagging (Fig. 2). As with other methods using GRMs based on array SNPs, LDAK-SC produced downwardly biased *h^2^_SNP_* estimates for rarer CV phenotypes. Using the MAF-stratified approach (LDAK-MS) resulted in similar patterns.

As with GREML-SC, using LDAK-SC on imputed data resulted in a complex set of biases that depended on CV MAF, data type, and stratification, although the patterns of bias were different. LDAK-SC *h^2^_SNP_* estimates using only common (MAF > 0.01) imputed variants were similar to those using only array SNP positions. LDAK-SC using all imputed variants led to roughly unbiased *h^2^_SNP_* estimates in unstratified samples, but led to *h^2^_SNP_* estimates that varied wildly depending on the CV MAF in the stratified samples (Fig. 2). LDAK-MS on imputed variants produced *h^2^_SNP_* estimates that were less biased that LDAK-SC, but nevertheless more biased and more sensitive to stratification compared to those produced by GREML-MS on imputed data (Fig. 2).

Using LDAK-SC on WGS data also resulted in biases. With only common variants, results mirrored those found using array and imputed variants (Fig. S5). However, when all WGS variants were used, *h^2^_SNP_* for very rare CV phenotypes was overestimated, especially in highly stratified samples, but underestimated for all other phenotypes. When using LDAK-MS on WGS data, the biases were less extreme. However, LDAK-MS resulted in over-estimated *h^2^_SNP_* for common CVs and underestimated *h^2^_SNP_* for all other CV phenotypes (Fig. 2). Similar to LDAK-MS estimates of total *h^2^_SNP_*, using LDAK-MS to partition genetic variance among MAF ranges, produced estimates that were less precise and more biased than either GREML-MS or GREML-LDMS for the array, imputed, or WGS based GRMs (Figs. 3, S7-S9, S12-S13).

Much of the observed patterns was likely due to the relationship between MAF and LDAK weights (Fig. S14; ref.^12^) and differences in MAF distributions of array, imputed, and WGS variants (Fig. S2). More very rare variants were observed and given higher weightings in the WGS data than in either the imputed or array datasets. Similarly, in stratified datasets more very rare variants were imputed (Fig. S2) and this likely contributed to stratification effects and differences among imputed and WGS datasets.

### Extended Genealogy with Thresholded GRMs

Patterns in the biases of *h^2^_SNP_* estimates were similar to those found using GREML-SC (Fig. 2) when using the extended genealogy method, demonstrating that *h^2^_SNP_* estimates are unaffected by the inclusion of close relatives so long as the model includes a second (thresholded) GRM that contains only information on genomic sharing among close relatives. However, the relative amount of variance attributable to the unthresholded GRM (estimating *h^2^_SNP_*) versus the thresholded GRM (estimating *h^2^_IBS>t_*) varied considerably, and depended on whether common (MAF>0.01; Fig. S15) or all (Fig. S16) markers were used to estimate the GRMs. Using GRMs built from common (MAF>.01) array markers (Fig. S15), the estimate of *h^2^_IBS>t_* was negative, while *h^2^_SNP_* was overestimated for common CV phenotypes. As CVs became rarer, *h^2^_IBS_>_t_* grew while *h^2^_SNP_* shrunk, consistent with Zaitlen et al.’s interpretation that *h^2^_IBS>t_* would estimate variance due to rarer CVs. This pattern was more pronounced when all markers were used (Fig. S16). Using imputed or WGS data, the pattern of negative variances estimated for some of the GRMs remained. Nevertheless, estimates of total heritability, similar to *h^2^_FAM_*, the sum of *h^2^_IBS>t_* and *h^2^_SNP_*, were nearly unbiased or slightly downwardly biased in most datasets and stratification subsamples (Fig. S15-S16). Even the total heritability of very rare CV phenotypes was underestimated by less than 5%, regardless of the dataset used (SNP, WGS, or imputed variant). It is important to note, however, that shared environmental effects can inflate estimates of total *h^2^* using this method (see *Confounding between relatedness and shared environments* below).

### Treelet Covariance Smoothing (TCS)

Estimates of *h^2^_SNP_* from the TCS approach were highly unstable. Using samples of unrelated individuals, the TCS method produced widely varying estimates of *h^2^_SNP_* depending on the CV MAF, level of stratification, and type of data used to build the GRM (Fig. 2). We note that the original implementation^21^ used related individuals for *h^2^_SNP_* estimatione however, performance did not improve when using samples of related individuals (Figs. S4-S6). The estimated and empirical standard errors were substantially higher than any other estimation method (Fig. S7-S9). Moreover, the pattern of results was complex and depended strongly on the simulation conditione for estimates from GRMs built from array (Fig. S4) or imputed (Fig. S6) markers, *h^2^_SNP_* was typically underestimated for all CV MAF frequencies irrespective of inclusion of close relatives. However, *h^2^_SNP_* estimates were too high when WGS data was used for certain combinations of CV MAF frequencies and stratification levels, and too low for others. It is possible the TCS method would work better in samples that included more close relatives, but it should be noted that other approaches (e.g., the thresholded GRM approach above) that rely upon inclusion of close relatives produced unbiased total estimates with our sample sizes.

### LD Score Regression

Estimates of *h^2^_SNP_* from LD Score Regression were similar when utilizing either Axiom SNPs, imputed, or WGS data (Figs. 2 and S4-S6), as were estimates of the intercept (which reflect the contribution of stratification and cryptic relatedness to the GWAS test statisticse Figs. S17-S19). Across data types, *h^2^_SNP_* was generally slightly underestimated (5-10%) for common CV phenotypes. This downward bias was slightly reduced in simulations using 10,000 causal variants, but remained (Fig. S20); it is possible that this bias would be eliminated under the truly infinitesimal model assumed by the model. *h^2^_SNP_* was increasingly underestimated for phenotypes caused by increasingly rare CVs (Fig. 2), regardless of data type. This underestimate of rare CV variation occurs because *h^2^_SNP_* is estimated only from common marker (MAF>0.01) GWAS statistics^22^, which are typically unaffected by rarer CVs. Interestingly, in the highly stratified subsample, common CV phenotype *h^2^_SNP_* was overestimated with no covariate correction with array SNPs, but controlling for PCs and sequencing cohorts using regression (Figs. 2 and S4-S6) or a mixedamodel approach (GCTA-LOCOe Fig. S4) removed this bias, suggesting that *h^2^_SNP_* estimates from LD score regression are not immune to biases due to stratification.

Estimates of *h^2^_SNP_* using MAF-partitioned LD score regression were highly variable, but in many cases biased upwards (Fig. S4-S6). For common CV phenotypes, the estimates were less biased than the standard LD score regression estimates described above. However, with rarer CV phenotypes, regardless of the data used (array positions, imputed variants, or WGS data), *h^2^_SNP_* was severely overestimated, expected when including very rare SNPs in the regression^11^.

The genomic control inflation factor, *λ*_GC_, was greater in more stratified subsamples without covariate correction, demonstrating the bias in GWAS with structure even in the absence of confounding environmental effects (Figs. S17-S19), consistent with previous work that shows structure alone can inflate GWAS test statistics^32,35,36^ due to chance CV allele frequency differences. We confirmed this using simulated data for two populations spanning a range of structure (*F_ST_*) and polygenicity without confounding environmental effects (Fig. S21). After controlling for PC covariates using regression or by inclusion of a kinship matrix (using GCTA-LOCO; Axiom SNPs only), there was limited effect of stratification, but *λ*_GC_ was still greater than one for phenotypes derived from common, uncommon and rare CVs (Figs. S17-S19). That *λ*_GC_ was not inflated for very rare CVs probably only reflects low statistical power for testing low MAF markers.

The LD score regression intercept, which reflects the amount of confounding by stratification and polygenicity^22^, was greater than one when no covariate control was applied across all stratification subsamples for all but the common CV traits (Figs. S17-S19). This was stronger for the more stratified subsamples, as expected. The intercept was ~1 when the covariates (and relatedness using Axiom SNPs) were accounted for, with the exception of uncommon and rare CV phenotypes, which were slightly >1, suggesting that the control of covariates was sufficient to account for the majority of the inflation in test statistics due to stratification. We note that these simulations included no confounding environmental effects, which may covary with stratification, and lead to inflation of GWAS statistics independent of the inflation of the intercept observed here^22^. Nevertheless, such inflated GWAS statistics generally should not be associated with the degree of LD-tagging of the markers, and thus should not inflate estimates of *h^2^_SNP_.*

### Confounding between relatedness and shared environments

We tested the effect of confounding between relatedness and shared environment (simulated *c*^2^= 0.1) for GREML-SC, LD score regression, and thresholded GRMs, using common array positions only. With unmodeled shared environmental effects, including all relatives and using GREML-SC resulted in overestimates of *h^2^_SNP_*, especially for rare CV phenotypes and for stratified samples (Fig. 4). However, when close relatives were removed at thresholds of 0.05 or 0.1, shared environmental effects produced no additional upward bias (Fig. 4) over those observed when no shared environmental effects existed (Fig. 2) Thus, as previously argued^6^ removing close relatives appears to correct for this type of shared environmental effect. Also as argued in ref.^22^, *h^2^_SNP_* from LD score regression was not biased upward due to unmodeled shared environmental effects, even when close relatives were included. Finally, using the thresholded GRM method with environmental confounding, *h^2^_SNP_* was biased slightly upward, particularly with a 0.1 relatedness threshold, but total heritability overestimation reached 20%, consistent with all or almost all shared environmental variance being estimated as additive genetic variance. Thus, care must be taken in interpreting results from methods that use SNP GRMs to estimate heritability when related individuals are included; shared environmental variance can masquerade as genetic variance.

### Heritability of Complex Traits in the UK Biobank

We applied the GREML-MS and GREML-LDMS approaches to six complex traits in the UK Biobank data, and partitioned estimates of the heritability by marker MAF using either directly genotyped Axiom SNPs or imputed genome-wide variants (Fig. 5, S23, Tables S6-S7). Total *h^2^_SNP_* was on average 12% lower using imputed data rather than the directly genotyped Axiom positions; our simulation results suggest this may be due to overestimation of variation due to common CVs for array markers (Fig. S4) and slight underestimation of variation due to common CVs for imputed markers (Fig. S6) using this method. The difference between array and imputed data was most apparent in the estimates of *h^2^_SNP_* per MAF bin, where *h^2^_SNP_* was lower using imputed data for common variant bins (MAF>0.01), but higher for rarer MAF bins. For example, the rare MAF bins (MAF<0.01) accounted for 8.8% of the phenotypic variance of height using imputed markers but only 0.6% using genotyped SNPs, whereas common MAF bins accounted for 48% and 59%, respectively. Fluid intelligence was even more striking, with rarer SNPs accounting for 11% and 3.4% using imputed and directly genotyped markers, respectively, while common markers accounted for 14% and 20%. Our simulations results suggest the *h^2^_SNP_* estimates from imputed data are more trustworthy.

**Figure 5.**
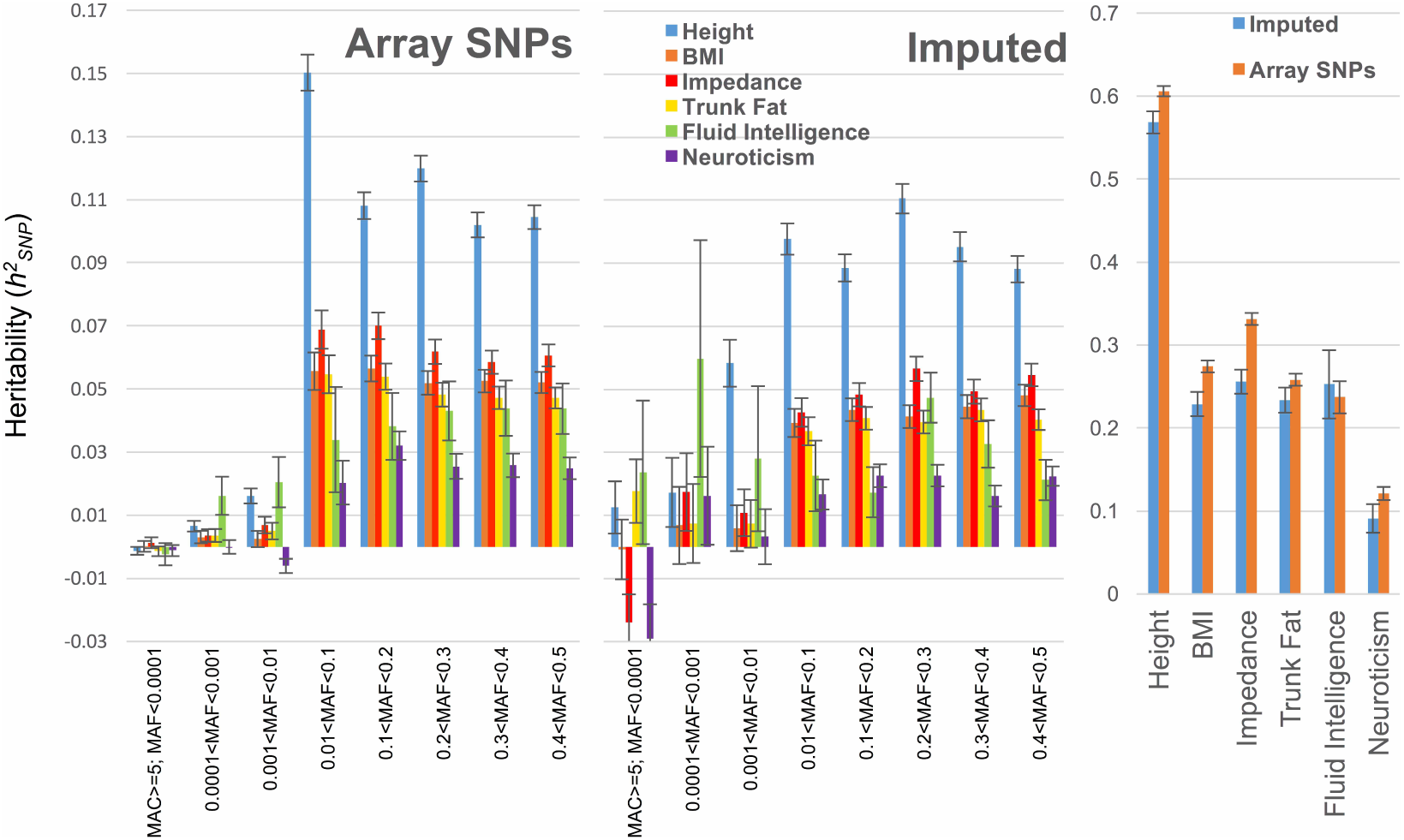
Estimates of MAF partitioned *h^2^_SNP_* using GREML-MS on Axiom array SNPs (left) and imputed genome-wide variants (center) for six complex traits in the UK Biobank. Total *h^2^_SNP_* shown on right.

Our simulation results also suggest that frequency distribution of CVs is best estimated using imputed data. The *h^2^_SNP_* across MAF bins from a GREML-MS model in the UK Biobank imputed data suggest real differences in genetic architectures across the six traits (Fig. 5). For example, height and adiposity phenotypes (BMI, impedance, and trunk fat) appear to be influenced mostly be common CVs, whereas fluid intelligence appears to have an important contribution from rare (MAF < 0.01) CVs. Results from GREML-MS (Fig. 5) were similar to those from GREML-LDMS (Fig. S23, Table S7), although GREML-LDMS suggested that more trait variance, even that attributable to common SNPs, was due to variants in the lower half of LD scores.

Our results suggest that most of the genetic variance of these traits is attributable to relatively common (MAF>0.01) variants. However, the contribution of increasingly rare CVs is likely to be underestimated for a few reasons. First, our simulations suggest that variation due to very rare CVs (0.0003<MAF<0.0025) is underestimated by ~ 20% due to low imputation quality of rarer variants. Second, this underestimate was probably more severe in these results given the imputation reference panel used in the UK Biobank data was half the size of the reference panel used in our simulations. The variation due to CVs not present in the imputation reference panel used for the UK Biobank (UK10K and 1,000 Genomes) were missed in our results.

## DISCUSSION

### Performance of *h^2^_SNP_* Methods in Simulated Data

We have demonstrated that estimates of genetic variation using SNP data can be biased in a number of sometimes difficult to foresee ways, and depend strongly on a complex interplay between method used, the frequency distribution of CVs, the type of data used in the analysis, the degree of sample stratification, whether relatives are included or excluded, and the importance of shared environmental effects. Approaches that are able to explore genetic architecture of complex traits also differ in their ability to correctly estimate the CV frequency distributions. Understanding how the different methods behave under different contexts is crucial for proper interpretation of SNP-heritability estimates and for optimal design of future studies. There has been much debate surrounding the relative importance of common vs. rare variants and the degree to which heritability remains unexplained (e.g., ref. ^7,12,13^), and the findings presented here offer context for how results from these methods inform these debates.

Through simulations, we have provided evidence that the use of WGS data, in combination with genome partitioning methods such as GREML-MS or GREML-LDMS, results in roughly unbiased *h^2^* estimates in unrelated samples, regardless of trait genetic architecture or population stratification in the sample, although variation due to extremely rare variants (e.g., de novo mutations) that are unshared between individuals in the sample will still be missed. Even with the most comprehensive imputation reference panel available, using imputed genome-wide markers still results in downwardly biased *h^2^_SNP_* estimates to the degree that rare variants are important to trait variation, but not nearly to the degree observed when using array markers. This is important, because it implies that the narrow-sense heritability remains underestimated in current studies using imputed data. Even with datasets using large reference panels, such as the UK Biobank data presented here, *h^2^_SNP_* from very rare CVs is likely underestimated due to poor imputation of rare SNPs. As imputation reference panels, such as the HRC and the forthcoming TOPMed panel^37^, continue to grow in size and diversity, accurate imputation of increasingly rarer variants will allow for increasingly accurate estimation of not the full narrow sense heritability, as well as for increasingly accurate estimation of the frequency distribution of CVs. Alternatively, novel methods, such as those that rely on sharing at identicalabyadescent haplotypes rather than allele sharing at measured SNPs^38^, may betteracapture effects of rare and poorlyatagged variants, and is a potential future direction for estimating the variation due to rare CVs.

Linkage disequilibrium (LD) between CVs and markers is central to the methods reviewed here. The observed patterns of over- and underestimation can be partly understood through the effect of LD among causal variants and markers (Fig. S11). As Yang et al^12^. demonstrated, using GREML-SC, *h^2^_SNP_* estimates should be unbiased when the average LD between markers and CVs (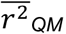) is the same as the average LD among all markers (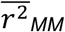), which occurs when markers and CVs are sampled from the same allele frequency distribution. This explains the underestimate of *h^2^_SNP_* using array genotypes when the CVs are rare, because common markers on an array typically have lower LD with rare CVs than with other markers, leading to (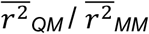) and *h^2^_SNP_* << *h^2^.* On the other hand, when the CVs are a random sample of markers, this ratio is ~1 and the estimated *h^2^_SNP_* ≈ *h^2^.* Finally, when the CVs are more common than markers used to create the GRM, LD between common CVs and markers will typically be higher than LD among markers, leading to 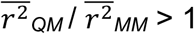 and *h^2^_SNP_* > *h^2^.*

The bias arising from a mismatch in CV and marker frequency distributions is not alleviated by weighting of markers by LD. Speed et al. showed that redundant marker tagging of CVs can bias *h^2^_SNP_* upward, and proposed weighting markers inversely to their LD score, which partially mitigates this bias in sparse genotype data. However, using such weights in dense whole genome sequence or imputed data leads to near 0 weights for most common markers, typically leading to underestimates of heritability arising from common CVs and, potentially, to overestimates of heritability from very rare CVs. What does appear to alleviate both the bias arising from a mismatch in CV and marker frequency distributions as well as the bias due to differential LD is binning markers by different MAF and LD bins^12^. When used on imputed or sequence data, GREML-MS and GREML-LDMS provide the most accurate partitioning of the variance and least biased total *h^2^_SNP_* estimates across genomic data types, CV frequency distributions, and levels of stratification. Although we showed that WGS is the ideal data source for creating GRMs, imputation will, for the time being, remain a cost-effective way to capture most of the trait variation, and will only improve as sequencing initiatives continue to amass larger, publicly available reference panels.

Our simulation results highlighted both limitations and advantages to LD score regression. Although it uses a much different approach than GREML, LD score regression suffers from many of the same problems as single-component GREML approaches. LD regression leverages the fact that for common variants under an infinitesimal model, the effect size of a marker is related to how well it tags the surrounding variants (and therefore how likely it is to tag a CV)^19,22^. Because LD is strongly related to MAF, the method increasingly underestimates variation as CVs become rarer. Moreover, unlike GREML-MS, it provides unreliable estimates if used on rare variants (MAF < .01), meaning that it cannot be used to accurately estimate CV frequency distributions, or variation due to rare CVs, even if GWAS statistics from imputed or WGS data are available. Nevertheless, LD score regression has several important advantages. Foremost among them, it can be used on summary statistics alone, bypassing the need for raw genotype data and allowing analyses based on sample sizes that would otherwise be impossible. Furthermore, as argued by its originators and as we have shown, it is generally robust to confounding biases due to stratification or shared family environmental effects, even when relatives are included in the sample. Finally, it is readily applied to various marker annotations in order to understand, for example, the relative importance of gene networks and functional categories^19^.

In our LD score regression simulation results, the contribution of common CVs to phenotypic variance were slightly underestimated, regardless of the data type used (array SNPs, imputed variants, or sequence data), a pattern previously reported^39^. This underestimate was not seen in the simulations performed by Bulik-Sullivan et al.^22^. This difference may stem from the fact that Bulik-Sullivan et al. simulated phenotypes caused by a much larger proportion of markers whereas we simulated phenotypes with only 1,000 or 10,000 CVs. Consistent with this possibility, when we increased the number of CVs to 10,000, our estimates were somewhat less biased. Nevertheless, it seems unlikely that the infinitesimal model truly holds for any phenotype, and thus *h^2^_SNP_* estimates from LD-regression are likely to be biased downward, especially as CVs become rarer.

There are several limitations to the findings presented here. First, although a subset of our simulations included shared environmental effects among close relatives, we did not model more complicated ways that environmental and genetic similarity can be confounded. For example, we did not simulate “vertical transmission” models in which distant ancestry can lead to low levels of environmental similarity, nor situations where environmental effects are confounded with ancestry. Previous studies have investigated this latter issue^40,41^, and fitting ancestry PCs removes much of the bias.

Second, other than varying CV MAF frequency distributions, we did not simulate situations where the LD of CVs differed systematically from the LD of markers used to estimate the GRM. As Speed et al. ^13^ demonstrated, if CVs come from regions of low LD (e.g., DNase I-hypersensitivity sites^42^), *h^2^_SNP_* will be underestimated and vice-versa when CVs come from regions of high LD. Yang et al.^12^ have shown that GREML-LDMS accounts for LD differences between CVs and markers and provides unbiased estimates. However, as we shown (Fig. S7-S9), standard errors for GREML-LDMS results are higher than GREML-MS. Given this tradeoff, we recommend that investigators report results from both approaches, and trust those from GREML-LDMS if there is a difference.

Third, we simulated CV effect sizes that were proportional to their minor allele frequencies (∝ [p(1ap)]^−α^, where a = a1 in nomenclature of ref. ^13^), so that the pe^variant contribution to heritability remained constant across MAF, similar to other studies^6,43,44^. The validity of this assumption has been the subject of recent debate (e.g., ref. ^13,45,46^) and it is clear that if this assumption is unmet in real data, using a single component model will bias estimates, as several well-designed evaluations of GREML-SC and LDAK-SC have shown^13^. However, two relevant findings from those studies bear mentioning. First, the scaling we applied (α = −1) is the most robust to violations of the model assumptions, and in sensitivity analyses of real data, scaling with various approaches often led to qualitatively and quantitatively similar conclusions^12^. Second, the GREML-MS and GREML-LDMS stratified approaches allow variances to differ across MAF partitions, effectively achieving the same goal as varying the scaling factor and allowing a greater exploration of CV frequency distributions. An interesting avenue of future work could be exploring possible values of α among functional annotations for evidence of purifying or positive selection.

### *h^2^_SNP_* Estimates in the UK Biobank

Using over 120,000 individuals with imputed genome-wide variants, we obtained estimates of *h^2^_SNP_* for complex traits similar to those previously published using directly genotyped markers and imputed genome wide markers for height and BMI (e.g., ref.^12^). Estimates of *h^2^_SNP_* for measures of adiposity (impedance, trunk fat, and BMI) were similar to each other, as expected given the relationship between these traits. Accounting for imperfect imputation and using our simulation results as guidance, our results suggest that the true narrow-sense heritability of height is 60-70%, and that of BMI is 20-30%, with some additional variation possibly from very rare and poorly-imputed CVs. Furthermore, the majority (~80%) of the additive genetic variance in these complex traits is explained by common variants with small additive effects, with a smaller proportion attributable to rarer variants. This finding has been discussed elsewhere^6,12,13^. This indicates that larger sample sizes will be required to identify common variants of very small effects in GWAS, but that little still-missing additive genetic variation remains.

The two behavioral traits we examined appear to have qualitatively different genetic architectures. Little of the additive genetic variance in neuroticism was explained by rare variants, but roughly half of fluid intelligence *h^2^_SNP_* was explained by rare variants with MAF < 0.01. Family- and twin-based estimates of heritability of intelligence are ~50%, while recent studies using common SNPs have estimated *h^2^_SNP_* ~ 0.25^47,48^. Our estimates, using an independent sample, are not dissimilar from these, and accounting for the downward bias in *h^2^_SNP_* using imputed data, heritability is likely ~30%, with roughly half of that from rare variants, and some additional variance caused by very rare and poorly-imputed CVs. However, given that we know that variation due to increasingly rare CVs is increasingly underestimated, it is possible that a larger proportion of the additive genetic variation in fluid intelligence is due to extremely rare CVs. Nevertheless, 30% is substantially lower than the ~50% estimates from family-based studies. However, it is also possible that these twin- and family-based estimates are overestimated, and that little remaining heritability will be explained by increasingly rare CVs. Our estimates of neuroticism heritability suggest that little of the variance is due to rare SNPs. In the UK Biobank data, our estimate of *h^2^_SNP_* (0.09) is slightly higher than some published estimates (*h^2^_SNP_* = 0.06[ref. ^49^]), but lower than a recent study using the same UK Biobank data (*h^2^_SNP_* = 0.14-16[ref. ^50^]). This may be due to our use of MAF-stratified GREML, rather than single component GREML with array data as in Smith et al., which we have showed here leads to overestimation of variance due to common CVs. Extended-twin family studies, which can provide estimates of narrow-sense heritability while addressing concerns of shared environmental and non-additive genetic influences, suggest that the narrow-sense heritability of neuroticism is ~30%^10^, which still leaves much of the additive genetic variance unexplained and presents a puzzle to be solved by future investigation.

### Conclusions

Heritability is a fundamental concept of genetics and its unbiased estimation is critical for understanding complex trait genetics as well as for designing better studies and obtaining a clearer picture of the possible explanatory power of GWAS. Below we provide our recommended best practices for studies aiming to estimate *h^2^_SNP_* and CV frequency distributions for complex traits. Even when applying these best approaches, heritability is still likely underestimated, but will improve as larger sample sizes, larger imputation panels, and better methods to account for rare variants are developed.

#### Recommended Practices

- Careful quality control in genetic data, for instance based on missingness and Hardy-Weinberg equilibrium, is critical, particularly for caseacontrol data and/or when the sample is comprised of multiple cohorts^44^.
- Include appropriate covariates, such as principal components, cohorts, and other potential confounders as fixed effects in GREML models and in the GWAS models for LD score regression.
- MAF- and/or LD-stratified GREML approaches^12^ on WGS or imputed data provide the most accurate estimates of *h^2^_SNP_* and CV frequency distributions. Even if CV frequency distributions are not of interest, these methods provide the most accurate estimates of *h^2^_SNP_* and are also the most robust to biases caused by stratification and differences between the CV and marker allele frequency distributions. However, there is a bias-precision tradeoff: more GRMs lead to larger standard errors, necessitating larger sample sizes for these methods. We recommend to report results from both GREML-LDMS and GREML-MS, and to trust the results of GREML-LDMS if there is a meaningful difference.
- If possible, run GREML models on WGS data if available, and otherwise data imputed using the largest and most diverse reference panel possible. Currently, this is the HRC.
- If raw genomic data is not available, use LD score regression on summary statistics, but calculate LD scores using a large sequence reference panel. Estimates from LD score regression are typically lower than those produced by GREML-SC on array data.
- Related individuals may share common environmental and non-additive genetic effects that can inflate estimates of *h^2^_SNP_.* Removing related individuals provides estimates that are less likely to be inflated by such environmental and non-additive genetic factors.
- Most reports of *h^2^_SNP_* in the literature have used the GREML-SC approach. However, as we have demonstrated, these estimates are subject to a number of sometimes conflicting biases, making interpretation of GREML-SC results challenging. Most crucially, GREML-SC is especially sensitive to the similarity between the frequency distributions of the CVs and the markers used to create the GRM, which can differ across genomic data types and array types. Moreover, GREML-SC can be sensitive to stratification effects, even when ancestry covariates are included in the model.

## SUPPLEMENTAL DATA DESCRIPTION

The supplemental data includes 22 additional figures and 7 additional tables.

## CONSORTIA

### Haplotype Reference Consortium

Shane McCarthy, Sayantan Das, Warren Kretzschmar, Olivier Delaneau, Andrew R Wood, Alexander Teumer, Hyun Min Kang, Christian Fuchsberger, Petr Danecek, Kevin Sharp, Yang Luo, Carlo Sidore, Alan Kwong, Nicholas Timpson, Seppo Koskinen, Scott Vrieze, Laura J Scott, He Zhang, Anubha Mahajan, Jan Veldink, Ulrike Peters, Carlos Pato, Cornelia M van Duijn, Christopher E Gillies, Ilaria Gandin, Massimo Mezzavilla, Arthur Gilly, Massimiliano Cocca, Michela Traglia, Andrea Angius, Jeffrey C Barrett, Dorrett Boomsma, Kari Branham, Gerome Breen, Chad M Brummett, Fabio Busonero, Harry Campbell, Andrew Chan, Sai Chen, Emily Chew, Francis S Collins, Laura J Corbin, George Davey Smith, George Dedoussis, Marcus Dorr, AlikiaEleni Farmaki, Luigi Ferrucci, Lukas Forer, Ross M Fraser, Stacey Gabriel, Shawn Levy, Leif Groop, Tabitha Harrison, Andrew Hattersley, Oddgeir L Holmen, Kristian Hveem, Matthias Kretzler, James C Lee, Matt McGue, Thomas Meitinger, David Melzer, Josine L Min, Karen L Mohlke, John B Vincent, Matthias Nauck, Deborah Nickerson, Aarno Palotie, Michele Pato, Nicola Pirastu, Melvin Mclnnis, J Brent Richards, Cinzia Sala, Veikko Salomaa, David Schlessinger, Sebastian Schoenherr, P Eline Slagboom, Kerrin Small, Timothy Spector, Dwight Stambolian, Marcus Tuke, Jaakko Tuomilehto, Leonard H Van den Berg, Wouter Van Rheenen, Uwe Volker, Cisca Wijmenga, Daniela Toniolo, Eleftheria Zeggini, Paolo Gasparini, Matthew G Sampson, James F Wilson, Timothy Frayling, Paul I W de Bakker, Morris A Swertz, Steven McCarroll, Charles Kooperberg, Annelot Dekker, David Altshuler, Cristen Willer, William Iacono, Samuli Ripatti, Nicole Soranzo, Klaudia Walter, Anand Swaroop, Francesco Cucca, Carl A Anderson, Richard M Myers, Michael Boehnke, Mark I McCarthy, Richard Durbin, Gongalo Abecasis, & Jonathan Marchini

## ACKNOWLEDGMENTS

This work was supported by NIH grant R01MH100141 (to MCK), NHMRC grants 1078037 (PMV) and 1113400 (PMV and JY), and Sylvia & Charles Viertel Charitable Foundation Senior Medical Research Fellowship (JY). We thank the participants of the individual HRC cohorts. This research has been conducted using the UK Biobank Resource. We thank Doug Speed for providing LDAK5. We thank the Keller and Vrieze lab groups, the Institute for Behavioral Genetics, the CU Research Computing facility, and Sean Caron.

## WEB RESOURCES

BOLT-REML: https://data.broadinstitute.org/alkesgroup/BOLT-LMM/

GCTA: http://cnsgenomics.com/software/gcta/index.html

Haplotype Reference Consortium: http://www.haplotypeareferenceaconsortium.org/home

LD score regression: github.com/bulik/ldsc/wiki

LDAK: http://dougspeed.com/ldak/

UK Biobank: http://www.ukbiobank.ac.uk/

